# Subtle Cellular Phenotypes Inform Pathological and Benign Genetic Mutants in the Iduronidase-2 Sulfatase Gene

**DOI:** 10.1101/2025.04.17.649412

**Authors:** Anushka Viswanathan, Serena Elia, Steven Q. Le, Sarah Hurt, Balraj Doray, Jason Waligorski, Kylan Kelley, William Buchser, Patricia Dickson

**Affiliations:** College of Arts & Sciences, Washington University in St. Louis, St. Louis, MO; Department of Genetics, Washington University School of Medicine, St. Louis, MO; Department of Pediatrics, Washington University School of Medicine, St. Louis, MO; Roy and Diana Vagelos Division of Biology & Biomedical Sciences, Washington University School of Medicine, St. Louis, MO; Department of Veterinary Pathobiology, College of Veterinary Medicine and Bond Life Sciences Center, University of Missouri, Columbia, MO; McDonnell Genome Institute, Washington University School of Medicine, St. Louis, MO

## Abstract

As with most enzyme-storage disorders, DNA testing at an early age is critical for identifying genetic variants and their impact on disease burden. Still, most variants in genes such as Iduronate-2-sulfatase, *IDS,* are Variants of Uncertain Significance (VUS). In patients presenting with Hunter Syndrome, clinical testing for IDS enzyme activity has been the mainstay to determine whether a variant is likely damaging. However, these enzyme assays are unable to predict disease severity or neuronal toxicity and may be missing many aspects of IDS pathology. In this study, we have developed an image-based assay using genome-engineered cells with *IDS* mutations to identify if a specific mutation causes lysosomal and membrane disruptions that characterize the disease. Specifically, we generate twelve mutant cell lines and document both the biochemical changes in IDS activity and the reproducible phenotypic differences therein. Next, we examine two patient-derived cell lines and find the same phenotypic differences in these lines compared to parental controls. The phenotypic changes are measured on a specific scale, which we term the PathScore_LC_. To determine whether the observed changes are specific to IDS, we reintroduce a recombinant version of the IDS enzyme (rhIDS) to rescue both the biochemical and phenotypic changes of these cells. Interestingly, even after 120 hours, rescue is present, but not all the cells return to normal, which may also reflect the pathological state. Finally, we examine the gene expression differences and find that recombinant enzyme is not sufficient to induce transcriptional changes in the mutant lines at the time points studied. Overall, these cell-based lysosomal and membrane phenotypes may be key to quickly and accurately profiling clinical variants in the IDS gene.

## Introduction

The classification of variants in the *IDS* gene through molecular DNA testing is hampered by difficulties in interpreting variant pathogenicity accurately, which complicates the differentiation between mucopolysaccharidosis type II (MPS II) phenotypes. To address these challenges, investigators have established standardized criteria, incorporating *in silico* prediction tools, population frequency data from resources such as gnomAD, and assessments of whether a variant is inherited or arose de novo in the proband.^1,2^ However, *in silico* models could potentially overestimate the functional impact of pathogenic variants, as even a small percent of IDS enzyme activity is corrective.^3^ In vitro assays which utilize artificial substrates present challenges in its own right, including the potential to misclassify pseudodeficiency alleles as pathogenic, resulting in false-positive findings.^4,5^ Despite these approaches with their limitations, a significant proportion of variants remain unclassified, often categorized as variants of uncertain significance (VUS). VUS have been reported in around 20% of genetic tests, highlighting their prevalence.^6^ Newborn screening for MPS II, while accepted into the Recommended Uniform Screening Panel (RUSP), is available in twelve states at the time of this writing.^7,8^ These obstacles in classification underscore discrepancies among researchers and clinicians regarding the interpretation and application of established criteria.^9,10^ Early detection is crucial, as individuals with MPS II exhibit early onset symptoms, reduced lifespans, and irreversible damage due to the progression of the disease.^11–13^ Thus, advancing predictive models holds promise for mitigating these challenges, ultimately enhancing diagnostic accuracy, improving patient care, and advancing our understanding of *IDS* gene variants.

Efforts to address challenges with variant classification, especially for VUS are ongoing. Platforms such as GeneMatcher and other voluntary reporting services assist by collating information on genetic variants along with phenotypic data to identify similar cases.^14,15^ *In silico* prediction tools, though imperfect, are still capable of rapidly evaluating nucleotide variants to determine their impact on protein function.^16–18^ This tool, however, relies heavily on a deep understanding of protein functional domains which still lacks completeness. The use of model organisms for studying genetic variants is promising, but is of low throughput and limited by reliance on orthologues and intronic variants that are different across species.^19^ Clinical whole exome sequencing (WES) or genome sequencing have become cost-effective approaches with high diagnostic yields, leading to broader use.^20–23^ The American College of Medical Genetics and Genomics (ACMG) now recommends it as a first- or second-line test for developmental delay or intellectual disability manifesting before age 18 years, or congenital anomalies with onset under age 1 year.^23^ However, WES and genome sequencing present their own challenges, particularly in interpreting the less well-characterized variants often seen in Hunter Syndrome. Cell-based assays utilizing patient-derived human cell lines exist but are hindered by a lack of functional assessments that are available to determine the pathogenicity of variants in a single cell.

We attempt to fulfill this need by using cellular phenotypes to proxy disease pathogenicity and aid variant classification. In this study, we tested our model’s ability and fitness to classify *IDS* genetic variants known to cause MPS II, also referred to as Hunter Syndrome. This X-linked disorder that is due to mutations in the *IDS* gene leads to reduced production of the lysosomal enzyme iduronate-2-sulfatase (I2S; referred to hereafter for simplicity as IDS), which is responsible for degrading the glycosaminoglycans (GAGs), heparan sulfate and dermatan sulfate, in the lysosome.^24^ In the absence of this critical enzyme, GAGs build up in the lysosome and cause systemic damage across many cells, tissues, and organs.^25^ Although biochemical assays such as IDS enzymatic assays and GAG measurements are available, a significant proportion of VUS necessitates a new methods of classification.^26^ We aim to address this by using image-based screening to differentiate between various classifications of variants in the *IDS* gene, with the goal of improving the categorization of VUS.

## Results

### Section 1: IDS Variants Affect both Enzyme Catalytic Ability and Cellular Phenotype

First, we chose a set of ‘anchor’ variants for *IDS* including known severe (pathogenic), known attenuated (pathogenic), and known benign missense variants (**Supplemental Table S1** and **Figure 1A**). We then engineered these variants into an A549 cell line (**Supplemental Table S2**). The A549 cell line has more than one X chromosome, so variants were present in combination. To evaluate the cellular impact of *IDS* mutations, we compared biochemical activity assays and pathogenicity scores based on a combination of phenotypes obtained from cellular imaging of these A549 cells, including morphological (shape) features, and lysosome and cell membrane intensity measures (**Supplemental Table S3**).

**Figure 1.**
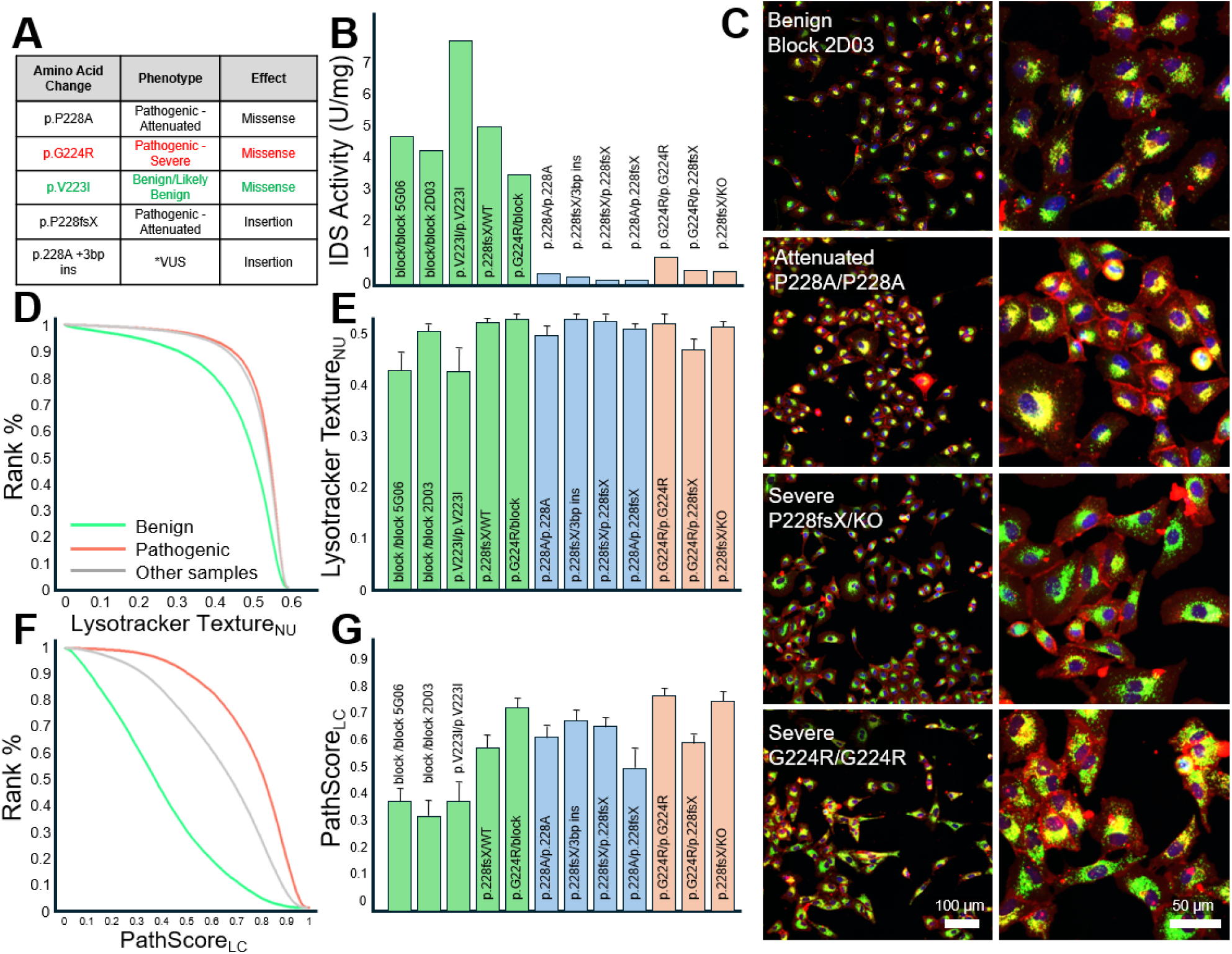
Phenotypic Signatures in A549 cells with Point Mutations in the *IDS* Gene. (**A**) Selected variants curated from ClinVar with documented clinical phenotypes and genetic consequence. Red text indicates a severe variant and green a benign variant. *VUS is the variant of uncertain significance tested in this study. (**B**) IDS enzyme activity measured in the 12 isogenic variant cell lines and controls by fluorometric assay with a 4-methylumbelliferyl substrate. Severe genotypes are shaded in red, attenuated in blue, and benign in green. (**C**) Fluorescent confocal micrographs of A549 cells, with nuclear stain in blue, cell membrane in red, and lysosome in green. The left panels show a larger field of cells, magnified for more detail in the panels to the right. Four of the mutant lines are shown, one per row. (**D**) Cumulative histograms of every individual cell from multiple replicates. X-axis shows the lysotracker non-uniformity (NU) texture, and the y-axis the rank percentile of every cell, where red lines are the G224R/G224R and the P228fsX/KO pathogenic lines, green are the block/block and V223I benign lines, and gray are all other variants. (**E**) Bar chart displays the average lysotracker texture NU with error bars representing standard deviation of 20-24 well replicates from multiple plates. (**F,G**) Same series as D,E, but reflecting the PathScore from the lysosomal and cell membrane (LC), a composite feature learned from the pathogenic and benign cells. (F) Cumulative histogram and (G) summary bar-chart with mean and standard deviation. Kruskal-Wallis p-value∼0, H-stat 17,869 among cells.

Of the twelve *IDS*-variant A549 isogenic cell lines, the pathogenic mutants exhibited markedly reduced IDS activity, with values ranging from 0 to 0.8 units/mg protein (**Figure 1B**). As expected, there was no clear separation between severe and attenuated pathogenic variants based on enzymatic activity. In contrast, benign variants demonstrated higher IDS activity, ranging from 3.5 and 7.7 units/mg protein.

We next performed fluorescent confocal imaging of the A549 *IDS*-mutant cells (**Figure 1C**). We found that benign variant cell lines and control cells exhibited well-defined nuclei, and high cell density, while attenuated pathogenic variant line P228A/P228A and severe pathogenic line P228fsX/KO showed increased irregularities by severity, greater overlap of the lysosomal and cell membrane markers, and reduced density compared with benign variant and control cell lines. Cells from the severe pathogenic line, G224R/G224R, displayed more disorganization, pronounced colocalization of markers, and the lowest cell density. Quantifying individual features of the cells, such as the intensity of the lysosomal stain or the size of the cell, did not generally discriminate between benign and pathogenic-mutant cells (not shown). The best single feature that could delineate the samples was a lysosomal ‘texture’ feature, that measures the patterns in the lysosomal staining (**Figure 1D, E**), but was still poor at discriminating individual mutant cells. Next, we used a composite metric derived from multiple features, termed the *PathScore_LC_*(**Figure 1F,G**).^27^ Neither the lysosome, cell membrane, or nuclei alone performed as well as a classifier which took both the lysosomes and cell membrane into account (**Supplemental Figure S1**). The phenotypic PathScore increased according to severity from benign to pathogenic variants and was approximately inverse of the IDS activity levels. Interestingly, some variants that had intact enzyme activity due to expression from a single normal allele (G224R/block and P228fsX/WT) displayed higher-than-expected PathScores, potentially indicating an effect of haploinsufficiency on the cellular phenotype. Overall, these findings demonstrate that mutations affect both IDS production and cellular phenotypes in a quantifiable and distinct manner (**Supplemental Figure S2**), allowing us to determine the degree of cellular damage caused by these *IDS* mutations.

### Section 2: Patient-derived Fibroblasts Phenocopy both Enzymatic and Phenotypic Signatures of Genome-Engineered lines

After establishing the cellular phenotype in engineered A549 cells, we examined IDS enzymatic activity in patient-derived fibroblasts with naturally occurring *IDS* variants. Cell line GM13203 had been obtained from a severely affected three year-old male who was hemizygous for a 1 bp insertion at nucleotide 208 in exon 2 of the *IDS* gene (c.208insC) resulting in a frameshift and premature stop codon (H70PfsX29). These cells had IDS activity of 0 units/mg protein, whereas cells from the unaffected carrier mother (GM01392) showed IDS activity of 39.9 units/mg, as expected (**Figure 2A**). Both cell lines were imaged, and their PathScore was measured exactly as with the A549 cells. The patient-derived cells from the affected child had a much higher PathScore both on average (**Figure 2B**), and when considering each individual cell (**Figure 2C**), compared with the unaffected parent. This result confirmed the robustness of the image-based score across cell lines.

**Figure 2.**
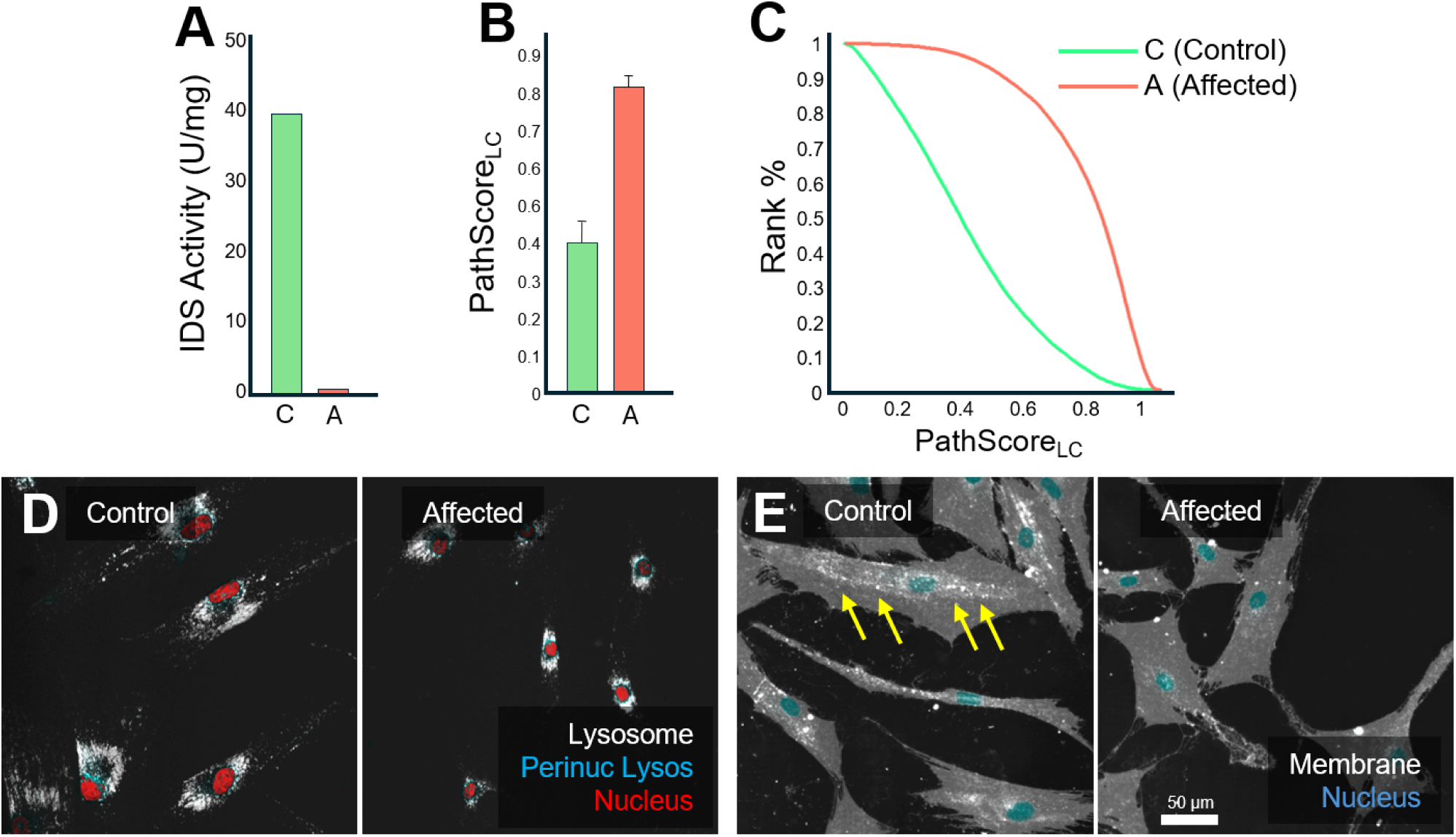
Cellular Morphology of Hunter Patient Fibroblasts. (**A**) IDS activity in patient fibroblasts of MPS II affected 3-year-old boy (GM13203) and the unaffected MPS I mother (GM01392). (**B**) Bar chart of PathScore_LC_ obtained from images of each patient fibroblast sample. Error bars are standard deviation. (**C**) Cumulative histogram of all cells from separate replicate wells shows the distribution of PathScores for the control and affected individuals. (**D**) Psuedo-color micrographs of fluorescent confocal images of fibroblasts with staining of lysosome (white), perinuclear lysosome (teal), and nucleus (red) between control and affected samples. (**E**) Staining of membrane (white) and nucleus (teal) in control and affected samples. Yellow arrows indicate a decrease presence of membrane inclusions found in mutant cells compared to control cells. Scalebar 50 µm.

Examining the fibroblasts more closely in terms of lysosomal staining (**Figure 2D**) and membrane staining (**Figure 2E**) revealed further insights. Lysosomes appeared similar in the two genotypes, although were spread subtly thinner throughout a large cytoplasm in control cells. The membrane in the affected cells showed decrease in intensity and a reduction in the number of membrane inclusions, suggesting a possible dysfunction in membrane turnover or recycling processes.^28^ These results imply poorer endocytosis and autophagic pathways in the mutants, which are critical for maintaining cell survival and homeostasis.

### Section 3: Recombinant Iduronate-2 Sulfatase enzyme restores wild-type Morphology in pathogenic Mutant A549 cell-lines

We next asked whether the addition of recombinant human IDS enzyme to *IDS*-mutant A549 cells would reverse the pathogenic phenotypes. One control (Block) and one benign variant (V223I/V223I) isogenic cell lines were compared to the attenuated (P228fsX/P228fsX) and severe (G224R/G224R) isogenic lines. First, we evaluated the IDS activity and confirmed a dose-dependent restoration of IDS activity by recombinant human IDS at 24 hours (**Figure 3A**). We next asked whether restoration of IDS enzyme activity by exogenous application of recombinant human IDS enzyme was sufficient to ‘correct’ the cellular phenotype. At the higher applied dose (concentration) of recombinant human IDS in the G224R/G224R variant cell line, a noticeable reduction in PathScore was observed when averaging cells among all time points after enzyme treatment (8 wells per condition, Kruskal-Wallis H-stat=26320, p∼0 **Figure 3B**). As anticipated, the cell-based PathScore_LC_ of enzyme-treated A549 cells showed no significant differences compared with the pre-treatment PathScore_LC_ for the benign variant and control cell lines. This finding demonstrates that treatment with enzyme may cause measurable corrections in cellular phenotype and suggests that at least some aspects of the cellular phenotype we observed may be caused by IDS deficiency.

**Figure 3:**
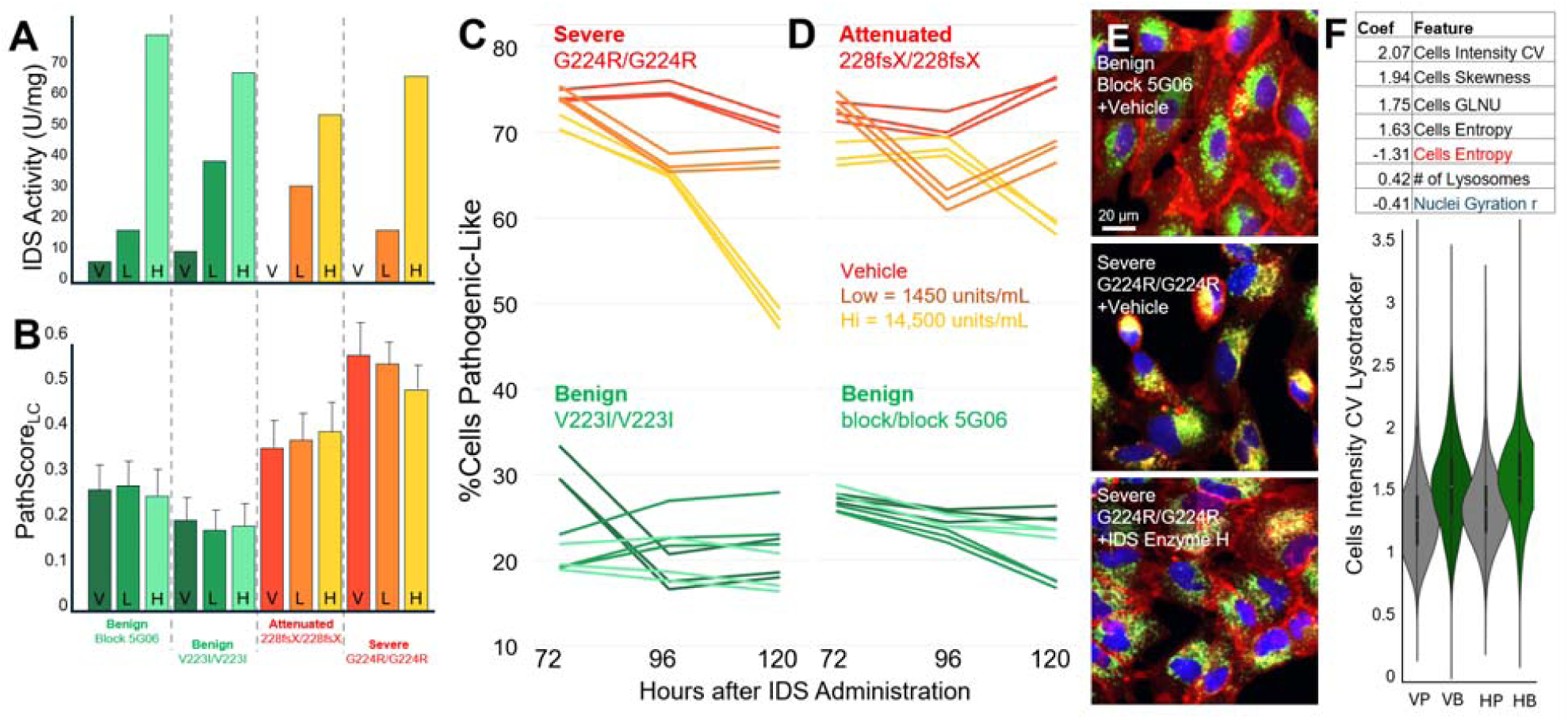
Recombinant human IDS restores partial wild-type phenotypes in A549 cells with severe and attenuated *IDS* variants. (**A**) Bar chart showing IDS enzyme activity of cell pellets for isogenic A549 lines with *IDS* variants of different severities, treated with a functional recombinant human IDS enzyme, 316.7 ug/mL = 14,471 units/ml (H), 31.67 ug/mL = 1,447 units/ml (L), or an artificial CSF vehicle (V) for 24 hours. One unit of activity = 1 nmol of converted substrate per hour. (**B**) Bar chart showing the deep cellular phenotype’s PathScore_LC_ for the same set of variants and conditions, averaging across replicate wells, plates, and time points. (**C, D**) Change in %cells pathogenic-like after rescued with recombinant IDS at 72, 96, and 120 hours after treatment. Top panels show change in severe variant (G224R/G224R, C and P228fsX/P228fsX, D) compared to benign variant (V223I/V223I, C and block/block, D) below. Notice that benign and mutant cells are non-overlapping on the y-axis, and that the high dose of IDS enzyme yields 50% normal cells after 120 hours. Brighter colors received the high IDS dose, darkest received vehicle, as shown in the bar graphs (**A, B**). Each line represents one of three plate replicates. (**E**) Example micrographs of treated cell lines with nuclei (blue), lysosomes (green), and cell membranes (red). The block-only benign control (top) is compared to the severe KO treated with the vehicle (middle) and a high dose of the IDS enzyme (bottom) at 120 hours. (**F**) Table listing features that contribute the most to whether high dose treated G224R mutant ‘responded.’ The negative signs indicate negative correlation such that higher values in this feature prevented a cell from responding. Black text is lysosomal features, red are cell membrane, and blue are nuclear features. Lower panel shows the top-ranked responsive feature (interaction term), and how it differs in cells that were vehicle-treated pathogenic (VP) or vehicle-treated benign (VB), and cells that were high dose-treated pathogenic (HP) or high dose-treated benign (HB).

While treatment with recombinant human IDS rapidly restores IDS activity in cells (typically within an hour or less after application), the catalysis of glycosaminoglycan substrate and subsequent restoration of the cellular phenotype may require more time. We next examined changes in the PathScore at varying time points several days after incubation with recombinant human IDS. The severe (G224R/G224R) variant cell line showed maximal PathScore_LC_ correction at 120 hours, with ∼50% of the cells displaying normal PathScore_LC_ at that timepoint (**Figure 3C**). The attenuated (P228fsX/P228fsX) variant cell line also showed correction at both the low and high doses (concentrations) of recombinant human IDS, with ∼40% of the cells displaying a normal PathScore_LC_ (**Figure 3D**).

Live-cell fluorescence imaging revealed a reduction in membrane staining and unipolar arrangement of the lysosomes in the severe (G224R/G224R) variant cells, consistent with the elevated PathScore_LC_ of this line (**Figure 3E**). We observed a partial restoration of the membrane and lysosome morphology following treatment with the high dose (concentration) of recombinant human IDS. We next examined the cellular features of the individual cells that were ‘rescued’ when treated with recombinant human IDS using a logistic regression model. The interaction terms that had the strongest significance (marking cells that had lower PathScores after treatment with rhIDS) were the cytoplasmic lysosomal intensity variation (Cell Intensity CV, coefficient = 2.07), followed by other lysosomal texture measurements (Cells Skewness,1.94, Cells GLNU, 1.75) (**Figure 3F**). As expected, when examining the lysosomal intensity variation across the population, it is the lowest in the ‘benign’-like cells (with a low PathScore_LC_) and increases in cells treated with the high dose of recombinant human IDS. Interestingly, only membrane stain texture (cells entropy) and nuclei shape (gyration radius) were negative coefficients, in that an increase in these metrics was associated with a more pathogenic phenotype, thereby negatively impacting phenotypic rescue. Together, these findings demonstrate that exogenous recombinant human IDS can rescue a portion of mutant cells’ cellular phenotypes, and that those changes are detectable via cellular imaging.

### Section 4 Transcriptional Signature is Distinct amongst *IDS* variant cell lines, and is not rescued by recombinant I2S enzyme after 120 Hours

Because restoration of IDS activity with exogenously applied recombinant human IDS did not completely rescue the cellular phenotype of *IDS* variant cell lines, we hypothesized that some aspects of the phenotype may be resistant to correction, and that these “irreversible” disease alterations may manifest in the cell’s gene expression profile. To investigate gene expression differences in Hunter syndrome (MPS II) cells and whether recombinant human IDS restores normal expression profiles, we performed bulk RNA sequencing on a subset of A549 mutant *IDS* cell lines, with and without recombinant human IDS treatment. Differential expression analysis identified several genes that were upregulated or downregulated in pathogenic genotypes compared to benign (**Figure 4A, B**). Expression of *IDS* was downregulated in affected cells compared to controls, as expected. An unsupervised UMAP clustering was performed on the RNA count table to determine, in an unbiased way, the underlying structure between the samples. The UMAP clearly separated the pathogenic and benign mutants, especially along axis 2 (**Figure 4C**). The mutant line was the strongest determinant of UMAP position, with time-of-treatment having a more modest effect. While rescue with recombinant human IDS led to improvements in biochemical assays and cellular phenotypic PathScore_LC_, the UMAP clustering indicated that treatment did not restore the transcriptomic profile to a healthy genetic state. This could indicate cell to cell differences acquired by the lines or indicate a broader impact of IDS beyond its immediate enzymatic activity.

**Figure 4.**
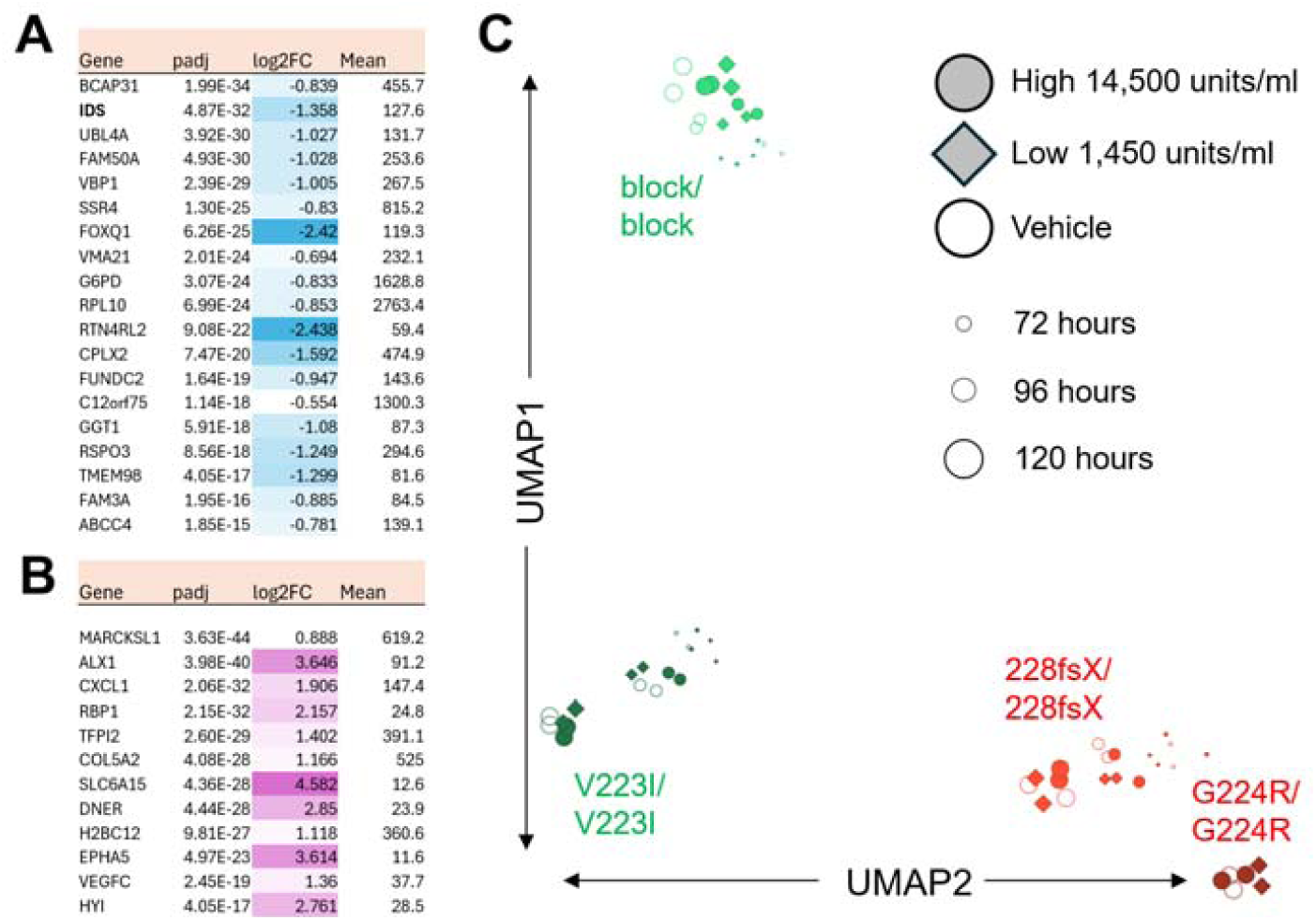
Transcriptional Profiling of *IDS* mutants and Recombinant Enzyme Treatment. (**A, B**) Heatmap-shaded table of differentially expressed genes in pathogenic (P228fsX/P228fsX, G224R/G224X) vs. benign (Block, V223I/V233I) conditions. All treatments and timepoints were pooled for these tables to maximize observations. Genes with both positive (upregulated, **A**) and negative (downregulated, **B**) log2 fold changes are listed from DeSeq2 analysis. Mean indicates the mean count of the gene across conditions. (**C**) UMAP visualization of transcriptomic profiles for selected variants after application of vehicle, low-dose recombinant human IDS, or high-dose recombinant human IDS, separated by time points. Filled markers are treated, while open markers were only vehicle exposed. Size of the marker indicates time after treatment, with largest markers indicating 120 hours.

## Discussion

In this study, we aimed to improve the classification of *IDS* gene variants, particularly those of uncertain significance (VUS), using cellular phenotyping. Our findings highlight the value of using cellular morphology and phenotypic data to aid in variant classification, thereby providing more nuanced insights into the pathogenicity of *IDS* mutations associated with MPS II.

The lysosomal accumulation of heparan sulfate and dermatan sulfate glycosaminoglycans occurs in Hunter syndrome and is thought to be the primary driver of the disease process. Abnormal accumulation of heparan sulfate in particular is detrimental to the central nervous system (CNS) and results in a wide range of clinical symptoms.^29^ Roughly two-thirds of individuals affected with Hunter syndrome have a clinically-evident neuronopathic disease and can experience developmental delays, cognitive decline, hyperactivity, seizures, behavioral challenges, and communicating hydrocephalus.^30,31^ Somatic manifestations, which are present in individuals who have non-neuronopathic as well as those with neuronopathic disease, include coarse facial features, growth delay, and pulmonary dysfunction.^30,31^ Newborn screening has been developed for Hunter syndrome, and twelve US states have active screening programs at present.^7,8,32^ There is a pressing need to develop methods that will predict which genotypes will result in the severe, neuronopathic form, because therapeutic options might one day be different for those children.

One of the major challenges in classifying *IDS* gene variants lies in the limitations of the biochemical assays for IDS activity and glycosaminoglycans, which may not reliably distinguish severe, neuronopathic from attenuated, non-neuronopathic phenotypes. In our small selection of *IDS* variants, we likewise found that the fluorometric IDS enzyme assay effectively differentiated the pathogenic variants from the benign variant and control cell lines. However, enzyme activity did not distinguish between the attenuated (P228fsX/P228fsX) variant line and the severe (G224R/G224R) variant line. By examining cellular phenotypes using our image-based assay, we found that the PathScore_LC_, a multi-feature indicator of cellular disruption, can be a reliable method for distinguishing between pathogenic and benign mutations, as well as varying levels of pathogenicity within the affected mutations (**Figure 1G**). This multi-dimensional approach, which captures cellular damage through the analysis of features such as cell size, lysosomal texture, and membrane integrity, provides a more complete picture of the molecular consequences of *IDS* mutations.

*In vitro* studies with A549 isogenic cell lines revealed that both severe and attenuated mutations exhibit significant morphological alterations, including changes in lysosomal and membrane distribution and cell shape (**Figure 1C**). These differences suggest possible disruptions in intracellular trafficking and cytoskeletal organization, which may underlie the pathogenic effects of severe mutations and are consistent with the MPS II disease state.^33^ These phenotypic changes were quantified using PathScores, revealing a clear gradient from benign to attenuated to pathogenic mutations (**Figure 1G**). This trend closely mirrored IDS enzyme activity levels (**Figure 1B**), further validating the utility of combining biochemical and phenotypic data in variant classification. Thus, the simultaneous use of multiple phenotypic features can greatly improve classification accuracy, providing a robust framework for distinguishing *IDS* variants.

Our analysis showed a phenotype in heterozygous cell lines, including G224R/block and P228fsX/WT, as reflected in the higher PathScores observed for these two variant lines compared to homozygous benign and control lines (**Figure 1G**). We are uncertain whether the cellular phenotype is caused by haploinsufficiency or a dominant, “gain-of-function” effect where the pathogenic mutations, G224R and P228fsX (**Figure 1A**), on one X chromosome may interfere with the function of the protein produced by the normal allele. This disruption can impact the cellular phenotype. In Hunter syndrome, skewed X-inactivation may result in affected females with one pathogenic IDS variant. We did not explore X-inactivation in our cell lines. However, another possible explanation for the intermediate phenotype seen in these heterozygous variant lines is that adult-onset neurodegenerative disorders have been increasingly recognized in otherwise normal carriers of autosomal recessive lysosomal storage disorders. For example, heterozygous carriers of Gaucher disease may develop Parkinson’s disease despite normal glucocerebrosidase enzyme activity, which is thought to be due to lysosomal dysfunction and impaired protein homeostasis.^34–36^ Similarly, heterozygous carriers of *NPC1* mutations may show late-onset neurodegeneration, even in the absence of classical Niemann-Pick type C disease pathology.^37–39^ Evidence of haploinsufficiency or dominant-negative effects in lysosomal proteins has been observed in other disorders. *Hexb* haploinsufficiency in an Alzheimer’s disease mouse model was shown to contribute to neurodegeneration, causing detectable brain changes and reinforcing the role of lysosomal dysfunction in disease pathology.^40^ Heterozygous mutations in Cathepsin F (CTSF) cause Type B Kufs disease, an adult-onset neuronal ceroid lipofuscinosis.^41^ Similarly, heterozygous pathogenic variants in sulfamidase (SGSH; the enzyme deficient in MPS IIIA) have been suggested as a risk factor for early-onset neurodegenerative disease, with observed phenotypic changes despite the fact that a substantial amount of enzyme activity remains.^36^ Together, these findings raise the possibility that even when enzyme activity is partially reduced or unaffected, subtle cellular disruptions due to dominant-negative effects or haploinsufficiency may contribute to disease pathology in detectable ways. While this effect is yet to be reported in Hunter syndrome, its presence in other lysosomal storage disorders and neurodegenerative diseases suggests a similar mechanism at work with these variants.

IDS activity and PathScores correlated well in patient fibroblasts, indicating that our model can effectively differentiate between control and affected cells (**Figure 2A-C**). Furthermore, our model may also help identify novel phenotypes in Hunter syndrome, as observed in the reduced intensity and number of membrane inclusions in affected cells (**Figure 2E**). While we expected to see an increase in the number, size, or intensity of membrane inclusions characteristic of a lysosomal storage disorder, we instead observed a reduction in these inclusions. This suggests a potential defect in membrane turnover or recycling. The lysosome has a myriad of functions, including degradation of membrane lipids, such as sphingolipids and cholesterol esters.^28,42^ Defects in membrane turnover can impact these processes, altering cellular function. Additionally, lysosomal distention can increase the size and number of membrane-contact sites with the endoplasmic reticulum, hindering lysosome movement along microtubules to peripheral locations.^43,44^ Other features, such as the reduction in intensity, number, and size of lysosomes and the nucleus, further suggest disruptions in membrane trafficking and degradation pathways (**Figure 2D-E**). These observations emphasize the potential of this model to drive new discoveries in Hunter syndrome.

In the recombinant human IDS rescue experiment with the severe variant (G224R/G224R) cell line, significant correction in PathScore was achieved when high concentrations of enzyme were added compared to vehicle and low concentrations (**Figure 3B**). The rescue effect was visible in all variants at 96 and 120 hours, with the most pronounced improvements seen in the high-dose treatment of the attenuated (228fsX/228fsX) and severe (G224R/G224R) variants (**Figure 3C-D**). No significant correction was observed in the attenuated variant cell line, which had a lesser PathScore compared to severe (G224R/G224R) cells. This supports the idea that sufficient enzyme supplementation over time can partially restore enzymatic function. Imaging and violin plots further illustrated this, as the severe (G224R/G224R) variant, after high-dose rhIDS treatment, regained some cellular phenotypes resembling the benign Block 2D03 vehicle condition, with increased cell number, intensity, and size (**Figure 3E-F).** These results indicate that recombinant IDS may show dose-dependent effects, and that the effects were most dramatic in the severe variant line, which had the most abnormal PathScore. Significant phenotypic improvements were primarily observed in the severe variant, suggesting that enzyme activity alone, or the lack thereof, may not fully account for cellular pathology. Our differential expression results from the RNA sequencing analysis confirm that even though loss of IDS disrupts gene regulation, recombinant enzyme treatment does not fully normalize gene expression (**Figure 4A**). This is further supported by UMAP clustering, which shows that IDS-rescued cells remain distinct from healthy controls, indicating that transcriptional recovery is incomplete. This misalignment between biochemical correction and transcriptomic recovery raises questions about the extent of cellular rescue and suggests that additional therapeutic strategies may be needed to fully restore cellular homeostasis in MPS II patients. Future studies could explore whether prolonged treatment or combination therapies can further normalize the transcriptome.

This study highlights the utility of integrating cellular phenotyping with biochemical assays to improve *IDS* variant classification in Hunter Syndrome. While enzyme activity assays distinguish pathogenic from benign variants, cellular phenotyping provides additional resolution, particularly for variants of uncertain significance. Additionally, rescue experiments demonstrated dose- and variant-dependent effects, revealing the potential for rescue treatments that can correct enzyme activity and some cellular features, especially in severe variants. These changes were detectable through imaging and quantified as PathScores using our model. However, our results show that despite improvements in IDS activity, the rescue effects remained limited, as the overall cellular phenotype remained disrupted. Building on these findings, future work may explore the Hunter syndrome cellular phenotypes in neurons to determine whether variants are associated with neuronopathic or non-neuronopathic forms of disease. These findings underscore the potential of image-based screening to improve diagnostic accuracy and inform treatment strategies for Hunter syndrome.

## Materials & Methods

### Variant Selection

Variants of the *IDS* gene were selected from ClinVar by a medical geneticist (P.I.D.) and sourced to publications in which the patient phenotype was comprehensively described. A total of twelve variants, ranging from benign to severe, were selected (**Supplemental Table 1**). These include the severe c.670G>C (p.G224R)^45^, attenuated c.673_674insC (p.P228fsX)^46,47^, and benign c.667G>A (p.V223I) variants. Additionally, a three-base pair insertion, resulting in the one codon addition of an alanine residue at position 228, was categorized as a VUS and examined in this study (**Figure 1A**). The Genome Engineering and Stem Cell Core at Washington University School of Medicine (GESC) generated gRNAs XCC415h.h.IDS.sp3 (GCTTATGATACCCAACGGCCNGG) and XCC415h.h.IDS.sp2 (GTGTGGCTTATGATACCCAANGG), along with a control single-stranded donor DNA (ssODN) incorporating only block modifications to prevent recutting for each variant. The twelve selected variants were generated in isogenic A549 cell lines. These cell lines are diploid-triploid and require mutations to be introduced into the *IDS* gene on one or both X chromosomes. The mutations were engineered using CRISPR-mediated homology-directed repair (HDR). The arrangement of the twelve variants, along with their genotypic details are listed in **Supplemental Table 1**.

### Cell Culture and Transfection

An *IDS-*myc-pcDNA construct containing engineered mutations was transfected into A549 cells developed by the GESC core. The resulting twelve isogenic cell lines were used as a training set to validate the model and the machine learning platform. The training set, comprising twelve variants, was transfected into female A549 cells that had undergone complete *IDS* knock-out and expressed a LAMP1-GFP reporter. Additionally, Hunter Syndrome human fibroblasts (GM01392 and GM13203) were transfected with this construct as well. These two patient fibroblast samples were procured from the Coriell Institute for Medical Research. These two patient fibroblast samples were obtained from the Coriell Institute for Medical Research. The control sample is from a 24-year-old mother (GM01392) who is a carrier of MPS I, also known as Hurler Syndrome, and has a normal genotype for MPS II. The affected sample is from a 3-year-old male (GM13203) who is hemizygous with a frameshift mutation in the *IDS* gene caused by a 1 bp insertion at nucleotide 208 in exon 2 (208insC). This mutation created a premature stop codon, consistent with the severe phenotype (H70PfsX29).

### IDS Protein Purification

In order to express and purify IDS from cell culture media, nucleotides encoding the anti-Protein C epitope (HPC4) were appended to the 3’ end of the cDNA, and in the process removing the nucleotides for the myc epitope. IDS-HPC4 and GNPT-S1S3 cDNA were co-transfected into Expi 293 cells growing in suspension as per the manufacturer’s protocol (Thermo Fisher). S1S3 phosphotransferase is a manufactured version of GlcNAc-1-phosphotransferase, a Golgi-complex enzyme that modifies lysosomal proteins with mannose 6-phosphate glycans for lysosomal targeting.^48^ We found that when expressed recombinantly, most of the secreted IDS fails to acquire the mannose 6-phosphate lysosomal targeting signal on their glycan chains, and that this was corrected with co-transfection of S1S3. The serum-free media containing the secreted IDS was collected 5 days post-transfection and purified on a HPC4-agarose matrix (Sigma-Millipore) according to the manufacturer’s protocol. Briefly, the equilibration and binding steps were performed using buffer containing 20 mM Tris-HCl pH 7.5, 100 mM NaCl, and 1mM CaCl_2_. The same buffer was used for the wash step except that the NaCl concentration was increased to 500 mM. The proteins were eluted in 1 ml fractions in buffer containing 20 mM Tris-HCl pH 7.5, 100 mM NaCl, and 5 mM EDTA. Protein purity was determined by SDS-PAGE and Coomassie staining of the gels.

### Cell Plating Assay and Confocal Imaging

Both the A549 lines engineered by GESC and Fibroblast lines were kept in an incubator at 37°C with 5% CO2 and monitored daily to assess growth and health. Lines were transferred to 12-well plates with media and were allowed to proliferate for 3-5 days. Trypsinized cells from the 12 well plates were then randomly arrayed (to reduce batch variation by averaging out potential plate-related effects) across two 96-well black clear bottom tissue culture treated plates using custom software and plated using a Biomek i5 liquid handling robot. The following day, plates with the randomized lines were first stained with SPY650 Nuclear dye (Cytoskeleton Inc., CY-SC501) at 1:1000, for one hour. Subsequently, the plates were stained with a master mix of Lysotracker Blue DND-22 (ThermoFisher Scientific, L7525), 50 nM, and Cell Mask Green (ThermoFisher Scientific, C37608), 1:2000, for 30 minutes. Plates were rinsed twice with respective medias and then immediately imaged on a high throughput content Molecular Devices ImageXpress HT.ai confocal microscope at 20x 0.45 NA utilizing live cell chamber capabilities (5% CO2, 37°C temperature). To optimize focus during imaging, each field of view overlapped by 8% of their area. Imaging settings were held constant throughout an experiment and were manually inspected to exclude any out-of-focus images. Images were analyzed with InCarta, filtered and compiled.

### Cellular Phenotype Analysis

To enable the platform to identify phenotypes of interest, the machine must first be trained using a suitable model. Training was conducted using the twelve selected variants detailed in **(Supplemental Table 1).** Tracing and feature extraction were performed on multiple sets of images obtained from repeated experiments using Cytiva’s INCarta.^27^ After features were extracted from the images, binary classification machine-learning models were developed that used quantitative phenotypic measurements from individual cells as response variables that defined each cell as wild type or pathogenic mutant. Three primary machine learning platforms were used, including Microsoft Azure ML, Tibco Spotfire, and TensorFlow to generate the described model.^27^ The model was specifically trained to recognize perturbed phenotypes solely based on the characteristics displayed in the training set.^27^

Single cells are picked at random, and their phenotypic features are iteratively combined to identify the combination of phenotypes that best differentiates wild-type cells from affected cells. **Supplemental Table 3** lists the morphological phenotypes examined by the platform. The data from cell features, imaging, and raft position mapping are integrated using custom software developed by the Buchser Lab.^27^ Each cell is assigned a quantitative value, referred to as the pathogenic score, which ranges from 0 to 1.0. The score is determined based on the combination of phenotypes it observes, with values near 1 indicating pathogenic mutant cells and those near 0 corresponding to the wild type cells. This image-based screening approach has been utilized by other groups.^49,50^

### IDS Enzyme Assay

Enzyme assays were conducted on isogenic cultures grown in 12-well plates. After trypsinization, between 100,000 and 300,000 cells were collected, pelleted, and lysed for assay. A 1.25 mmol/L solution of 4-methylumbelliferyl-α-L-iduronide 2-sulfate (4-MU-α-IdoA 2-sulfate, BIOSYNTH, Louisville, KY Cat# EM0301) was prepared by dissolving 2 mg of 4-MU-α-IdoA 2-sulfate in 3.36 mL buffer consisting of 0.1 M sodium acetate buffer pH 5.0, 10 mM lead acetate, and 0.02% sodium azide. The reaction was initiated by mixing 10 μL of each sample with 20 μL of the 1.25 mM 4-MU-α-IdoA 2-sulfate solution. The reaction mixture was then left to incubate for 4 hours at 37 °C in a water bath. This allowed any enzyme present in the samples to cleave the sulfate group from iduronide 2-sulfate from the 4-MU-α-IdoA 2-sulfate substrate. After the incubation period, the reaction was terminated using 40 μL of McIlvain buffer (0.4 M Na-phosphate/0.2 M citrate pH 4.5). The samples were then vortexed and centrifuged to form pellets. A second enzymatic reaction was initiated by adding 10 μL of partially purified recombinant alpha-L-iduronidase (IDU from R&D Systems) to each sample, followed by incubation at 37 °C in a water bath for approximately 24 hours. During this step, IDU cleaved the iduronide moiety, releasing 4-MU, which generated fluorescence detectable by a plate reader. The reaction was then stopped with 200 μL of glycine carbonate at pH 10.7.

Fluorescence was measured using a BioTek Synergy plate reader, with excitation and emission wavelengths set to 360 nm and 455 nm, respectively. Fluorescence values were normalized against a standard curve generated from 4-MU solutions with known concentrations to quantify the enzymatic activity of iduronate-2-sulfatase (IDS) in each sample. Enzymatic activity was expressed as units, with one unit defined as the conversion of 1 nmol of substrate per hour. The protocol for IDS enzyme assays was adapted from the method described by Voznyi et al. in their 2001 publication, “A fluorometric enzyme assay for the diagnosis of MPS II (Hunter disease).”^4^ The described methodology was applied to measure IDS activity in the 12 A549 cell line variants, patient-derived fibroblasts, and recombinant human IDS rescue experiments (see Methods: Cell Culture and Transfection, Uptake for Rescue Experiments).

### IDS Uptake for Rescue Experiments

Lysosomal enzymes, such as iduronate-2-sulfatase (IDS), are internalized by cells after binding to the mannose 6-phosphate receptor (M6PR) located on the plasma membrane.^51–53^ Purified recombinant human IDS, with a specific activity 45.8k Units/mg protein, was added to the cell culture media at two concentrations: 1,447 units/ml (“Low”) and 10x higher at 14,471 units/ml (“Hi”). Cells were incubated for 24 hours to allow for glycosaminoglycan (GAG) storage accumulation, enzyme uptake, and phenotype manifestation. Cells were then removed at varying time points, 72, 96, and 120 hours, and IDS enzymatic activity was assessed using the IDS activity assay described above. To eliminate any interference from lysosomal enzymes present in the serum, all cells were cultured with heat-inactivated serum to inactivate any external lysosomal enzymes. Cells were then imaged and analysed following methods outlined above. The percent of cells classified as pathogenic-like by the machine learning platform after this rescue was measured to see if the phenotype was corrected and to what extent as described in *Machine learning and Model Generation*.

### RNA-Sequencing (BRB-seq)

Bulk RNA barcoding and sequencing (BRB-seq) uses early-stage multiplexing and sample barcoding to produce 3’ cDNA libraries.^54^ BRB-seq multiplexed libraries were prepared using the Mercurius BRB-seq kit (Alithea Genomics). For each sample, a minimum of 100,000 cells was collected and lysed, and total RNA was extracted using the MagMAX mirVana Total RNA Isolation Kit (Thermo Scientific, Waltham, MA) according to the standard protocol. RNA concentration was measured using the Quant-it RNA HS Kit (Thermo Scientific) and then normalized to a total input of 500 ng. Libraries were sequenced by GTAC@MGI and demultiplexed counts were returned.

## Acknowledgements

We would like to acknowledge work from Lina Ali and, Samah Nour in the Buchser lab for their work on initial imaging assay development and working with the A549 and patient Fibroblasts. We would like to thank Christina Gurnett and her lab for the initial access to the A549 cell line and early experimentation. We thank the Genome Engineering and Stem Cell Center GESC at the Washington University in St. Louis for cell line engineering services. RNA sequencing was performed with the BRB-seq Pipeline thanks to Rob Mitra, Emma Gries, and Brian Muegge. Finally, we would like to acknowledge the patients and their families.

## References

1. Richards S, Aziz N, Bale S, Bick D, Das S, Gastier-Foster J, Grody WW, Hegde M, Lyon E, Spector E, Voelkerding K, Rehm HL; ACMG Laboratory Quality Assurance Committee. Standards and guidelines for the interpretation of sequence variants: a joint consensus recommendation of the American College of Medical Genetics and Genomics and the Association for Molecular Pathology. Genet Med. 2015 May;17(5):405–24. doi: 10.1038/gim.2015.30. Epub 2015 Mar 5. PMID: 25741868; PMCID: PMC4544753.

2. Gudmundsson, S. et al. Variant interpretation using population databases: Lessons from gnomAD. Hum Mutat (2021) doi:10.1002/HUMU.24309.

3. Stapleton M, Kubaski F, Mason RW, Yabe H, Suzuki Y, Orii KE, Orii T, Tomatsu S. Presentation and Treatments for Mucopolysaccharidosis Type II (MPS II; Hunter Syndrome). Expert Opin Orphan Drugs. 2017;5(4):295–307. doi: 10.1080/21678707.2017.1296761. Epub 2017 Mar 8. PMID: 29158997; PMCID: PMC5693349.

4. Voznyi YV, Keulemans JL, van Diggelen OP. A fluorimetric enzyme assay for the diagnosis of MPS II (Hunter disease). J Inherit Metab Dis. 2001 Nov;24(6):675–80. doi: 10.1023/a:1012763026526. PMID: 11768586.

5. Mucopolysaccharidosis type II | Newborn Screening. (n.d.). Newbornscreening.hrsa.gov. https://newbornscreening.hrsa.gov/conditions/mucopolysaccharidosis-type-ii

6. Rehm HL, Alaimo JT, Aradhya S, et al. The landscape of reported VUS in multi-gene panel and genomic testing: Time for a change. Genet Med. 2023;25(12):100947. doi:10.1016/j.gim.2023.100947

7. Baby’s First Test. (n.d.). Newborn screening education and resources. Genetic Alliance. https://www.babysfirsttest.org/

8. Millington DS, Ficicioglu C. Addition of MPS-II to the Recommended Uniform Screening Panel in the United States. Int J Neonatal Screen. 2022 Oct 11;8(4):55. doi: 10.3390/ijns8040055. PMID: 36278625; PMCID: PMC9624303.

9. Amendola, L. M. et al. Performance of ACMG-AMP Variant-Interpretation Guidelines among Nine Laboratories in the Clinical Sequencing Exploratory Research Consortium. Am J Hum Genet 98, 1067–1076 (2016).

10. Amendola, L. M. et al. Variant Classification Concordance using the ACMG-AMP Variant Interpretation Guidelines across Nine Genomic Implementation Research Studies. Am J Hum Genet 107, 932–941 (2020).

11. Grant N, Sohn YB, Ellinwood NM, Okenfuss E, Mendelsohn BA, Lynch LE, Braunlin EA, Harmatz PR, Eisengart JB. Timing is everything: Clinical courses of Hunter syndrome associated with age at initiation of therapy in a sibling pair. Mol Genet Metab Rep. 2022 Feb 2;30:100845. doi: 10.1016/j.ymgmr.2022.100845. PMID: 35242576; PMCID: PMC8856919.

12. Tomanin R, Zanetti A, Zaccariotto E, D’Avanzo F, Bellettato CM, Scarpa M. Gene therapy approaches for lysosomal storage disorders, a good model for the treatment of mendelian diseases. Acta Paediatr. 2012 Jul;101(7):692–701. doi: 10.1111/j.1651-2227.2012.02674.x. Epub 2012 Apr 11. PMID: 22428546.

13. Lin HY, Tu RY, Chern SR, Lo YT, Fran S, Wei FJ, Huang SF, Tsai SY, Chang YH, Lee CL, Lin SP, Chuang CK. Identification and Functional Characterization of *IDS* Gene Mutations Underlying Taiwanese Hunter Syndrome (Mucopolysaccharidosis Type II). Int J Mol Sci. 2019 Dec 23;21(1):114. doi: 10.3390/ijms21010114. PMID: 31877959; PMCID: PMC6982257.

14. Landrum, M. J. et al. ClinVar: Improving access to variant interpretations and supporting evidence. Nucleic Acids Research 46, D1062–D1067 (2018).

15. Boycott, K. M., Azzariti, D. R., Hamosh, A. & Rehm, H. L. Seven years since the launch of the Matchmaker Exchange: The evolution of genomic matchmaking. Hum Mutat 43, (2022).

16. Schwarz, J. M., Rödelsperger, C., Schuelke, M. & Seelow, D. MutationTaster evaluates disease-causing potential of sequence alterations. Nat Methods 7, 575–576 (2010).

17. Adzhubei, I. A. et al. A method and server for predicting damaging missense mutations. Nat Methods 7, 248–249 (2010).

18. Ng, P. C. & Henikoff, S. SIFT: Predicting amino acid changes that affect protein function. Nucleic Acids Res 31, 3812–3814 (2003).

19. Baldridge, D. et al. Model organisms contribute to diagnosis and discovery in the undiagnosed diseases network: Current state and a future vision. Orphanet Journal of Rare Diseases 16, 1–17 (2021).

20. Baldridge, D. et al. The Exome Clinic and the role of medical genetics expertise in the interpretation of exome sequencing results. Genetics in Medicine 19, 1040–1048 (2017).

21. Phillips, K. A. et al. US private payers’ perspectives on insurance coverage for genome sequencing versus exome sequencing: A study by the Clinical Sequencing Evidence-Generating Research Consortium (CSER). Genet Med 24, 238–244 (2022).

22. Trosman, J. R. et al. Perspectives of US private payers on insurance coverage for pediatric and prenatal exome sequencing: Results of a study from the Program in Prenatal and Pediatric Genomic Sequencing (P3EGS). Genet Med 22, 283–291 (2020).

23. Manickam, K. et al. Exome and genome sequencing for pediatric patients with congenital anomalies or intellectual disability: an evidence-based clinical guideline of the American College of Medical Genetics and Genomics (ACMG). Genet Med 23, 2029–2037 (2021).

24. Meikle, P. J., Fuller, M., & Hopwood, J. J. (2005). Lysosomal Degradation of Heparin and Heparan Sulfate. Elsevier EBooks, 285–311. 10.1016/b978-008044859-6/50011-3

25. Hashmi, M. S., & Gupta, V. (2020). Mucopolysaccharidosis Type II. PubMed; StatPearls Publishing. https://www.ncbi.nlm.nih.gov/books/NBK560829/

26. Agrawal, N. et al. Genotype-phenotype spectrum of 130 unrelated Indian families with Mucopolysaccharidosis type II. Eur J Med Genet 65, (2022).

27. Yenkin, A. L., Bramley, J. C., Kremitzki, C. L., Waligorski, J. E., Liebeskind, M. J., Xu, X. E., Chandrasekaran, V. D., Vakaki, M. A., Bachman, G. W., Mitra, R. D., Milbrandt, J. D., & Buchser, W. J. (2022). Pooled image-base screening of mitochondria with microraft isolation distinguishes pathogenic mitofusin 2 mutations. Communications Biology, 5(1), 1–14. 10.1038/s42003-022-04089-y

28. Tanaka, M., Kikuchi, T., Uno, H. et al. Turnover and flow of the cell membrane for cell migration. Sci Rep 7, 12970 (2017). 10.1038/s41598-017-13438-5

29. Minami K, Morimoto H, Morioka H, Imakiire A, Kinoshita M, Yamamoto R, Hirato T, Sonoda H. Pathogenic Roles of Heparan Sulfate and Its Use as a Biomarker in Mucopolysaccharidoses. Int J Mol Sci. 2022 Oct 3;23(19):11724. doi: 10.3390/ijms231911724. PMID: 36233030; PMCID: PMC9570396.

30. Wraith JE, Beck M, Giugliani R, Clarke J, Martin R, Muenzer J; HOS Investigators. Initial report from the Hunter Outcome Survey. Genet Med. 2008 Jul;10(7):508–16. doi: 10.1097/gim.0b013e31817701e6. PMID: 18580692.

31. Lau H, Harmatz P, Botha J, Audi J, Link B. Clinical characteristics and somatic burden of patients with mucopolysaccharidosis II with or without neurological involvement: An analysis from the Hunter Outcome Survey. Mol Genet Metab Rep. 2023 Sep 8;37:101005. doi: 10.1016/j.ymgmr.2023.101005. PMID: 38053935; PMCID: PMC10694755.

32. Ream MA, Lam WKK, Grosse SD, Ojodu J, Jones E, Prosser LA, Rosé AM, Comeau AM, Tanksley S, Powell CM, Kemper AR. Evidence and recommendation for mucopolysaccharidosis type II newborn screening in the United States. Genet Med. 2023 Feb;25(2):100330. doi: 10.1016/j.gim.2022.10.012. Epub 2022 Nov 29. PMID: 36445366; PMCID: PMC9905270.

33. Hampe CS, Yund BD, Orchard PJ, Lund TC, Wesley J, McIvor RS. Differences in MPS I and MPS II Disease Manifestations. Int J Mol Sci. 2021 Jul 23;22(15):7888. doi: 10.3390/ijms22157888. PMID: 34360653; PMCID: PMC8345985.

34. Sidransky E, Nalls MA, Aasly JO, Aharon-Peretz J, Annesi G, Barbosa ER, Bar-Shira A, Berg D, Bras J, Brice A, Chen CM, Clark LN, Condroyer C, De Marco EV, Dürr A, Eblan MJ, Fahn S, Farrer MJ, Fung HC, Gan-Or Z, Gasser T, Gershoni-Baruch R, Giladi N, Griffith A, Gurevich T, Januario C, Kropp P, Lang AE, Lee-Chen GJ, Lesage S, Marder K, Mata IF, Mirelman A, Mitsui J, Mizuta I, Nicoletti G, Oliveira C, Ottman R, Orr-Urtreger A, Pereira LV, Quattrone A, Rogaeva E, Rolfs A, Rosenbaum H, Rozenberg R, Samii A, Samaddar T, Schulte C, Sharma M, Singleton A, Spitz M, Tan EK, Tayebi N, Toda T, Troiano AR, Tsuji S, Wittstock M, Wolfsberg TG, Wu YR, Zabetian CP, Zhao Y, Ziegler SG. Multicenter analysis of glucocerebrosidase mutations in Parkinson’s disease. N Engl J Med. 2009 Oct 22;361(17):1651–61. doi: 10.1056/NEJMoa0901281. PMID: 19846850; PMCID: PMC2856322.

35. Robak LA, Jansen IE, van Rooij J, Uitterlinden AG, Kraaij R, Jankovic J; International Parkinson’s Disease Genomics Consortium (IPDGC); Heutink P, Shulman JM. Excessive burden of lysosomal storage disorder gene variants in Parkinson’s disease. Brain. 2017 Dec 1;140(12):3191–3203. doi: 10.1093/brain/awx285. PMID: 29140481; PMCID: PMC5841393.

36. Douglass ML, Beard H, Shoubridge A, Nazri N, King B, Trim PJ, Duplock SK, Snel MF, Hopwood JJ, Hemsley KM. Is SGSH heterozygosity a risk factor for early-onset neurodegenerative disease? J Inherit Metab Dis. 2021 May;44(3):763–776. doi: 10.1002/jimd.12359. Epub 2021 Jan 25. PMID: 33423317.

37. Benussi A, Cotelli MS, Cantoni V, Bertasi V, Turla M, Dardis A, Biasizzo J, Manenti R, Cotelli M, Padovani A, Borroni B. Clinical and neurophysiological characteristics of heterozygous *NPC1* carriers. JIMD Rep. 2019 Jun 28;49(1):80–88. doi: 10.1002/jmd2.12059. PMID: 31497485; PMCID: PMC6718120.

38. Lopergolo D, Bianchi S, Gallus GN, et al Familial Alzheimer’s disease associated with heterozygous *NPC1* mutation Journal of Medical Genetics 2024;61:332–339.

39. Schneider SA, Tahirovic S, Hardy J, Strupp M, Bremova-Ertl T. Do heterozygous mutations of Niemann-Pick type C predispose to late-onset neurodegeneration: a review of the literature. J Neurol. 2021 Jun;268(6):2055–2064. doi: 10.1007/s00415-019-09621-5. Epub 2019 Nov 7. PMID: 31701332.

40. Whyte LS, Fourrier C, Hassiotis S, Lau AA, Trim PJ, Hein LK, Hattersley KJ, Bensalem J, Hopwood JJ, Hemsley KM, Sargeant TJ. Lysosomal gene *Hexb* displays haploinsufficiency in a knock-in mouse model of Alzheimer’s disease. IBRO Neurosci Rep. 2022 Jan 20;12:131–141. doi: 10.1016/j.ibneur.2022.01.004. PMID: 35146484; PMCID: PMC8819126.

41. Smith KR, Dahl HH, Canafoglia L, Andermann E, Damiano J, Morbin M, Bruni AC, Giaccone G, Cossette P, Saftig P, Grötzinger J, Schwake M, Andermann F, Staropoli JF, Sims KB, Mole SE, Franceschetti S, Alexander NA, Cooper JD, Chapman HA, Carpenter S, Berkovic SF, Bahlo M. Cathepsin F mutations cause Type B Kufs disease, an adult-onset neuronal ceroid lipofuscinosis. Hum Mol Genet. 2013 Apr 1;22(7):1417–23. doi: 10.1093/hmg/dds558. Epub 2013 Jan 7. PMID: 23297359; PMCID: PMC3596852.

42. Thelen AM, Zoncu R. Emerging Roles for the Lysosome in Lipid Metabolism. Trends Cell Biol. 2017 Nov;27(11):833–850. doi: 10.1016/j.tcb.2017.07.006. Epub 2017 Aug 30. PMID: 28838620; PMCID: PMC5653458.

43. G. Scerra, V. De Pasquale, M. Scarcella, M. G. Caporaso, L. M. Pavone, and M. D’Agostino, “Lysosomal positioning diseases: beyond substrate storage,” Open Biology, vol. 12, no. 10, Oct. 2022.

44. Cabukusta B, Neefjes J. Mechanisms of lysosomal positioning and movement. Traffic. 2018 Oct;19(10):761–769. doi: 10.1111/tra.12587. Epub 2018 Jul 17. PMID: 29900632; PMCID: PMC6175085.

45. Kosuga, M. et al. Molecular diagnosis of 65 families with mucopolysaccharidosis type II (Hunter syndrome) characterized by 16 novel mutations in the IDS gene: Genetic, pathological, and structural studies on iduronate-2-sulfatase. Molecular Genetics and Metabolism 118, 190–197 (2016).

46. Sohn, Y. B. et al. Identification of 11 novel mutations in 49 Korean patients with mucopolysaccharidosis type II. Clinical Genetics 81, 185–190 (2012).

47. Kim, C. H. et al. Mutational spectrum of the iduronate 2 sulfatase gene in 25 unrelated Korean Hunter syndrome patients: Identification of 13 novel mutations. Human Mutation 21, 449–450 (2003).

48. Liu L, Lee WS, Doray B, Kornfeld S. Engineering of GlcNAc-1-Phosphotransferase for Production of Highly Phosphorylated Lysosomal Enzymes for Enzyme Replacement Therapy. Mol Ther Methods Clin Dev. 2017 Mar 29;5:59–65. doi: 10.1016/j.omtm.2017.03.006. PMID: 28480305; PMCID: PMC5415318.

49. Hasle, N. et al. High-throughput, microscope-based sorting to dissect cellular heterogeneity. Mol Syst Biol 16, (2020).

50. Feldman, D. et al. Optical Pooled Screens in Human Cells. Cell 179, 787–799.e17 (2019).

51. Bohnsack RN, Misra SK, Liu J, Ishihara-Aoki M, Pereckas M, Aoki K, Ren G, Sharp JS, Dahms NM. Lysosomal enzyme binding to the cation-independent mannose 6-phosphate receptor is regulated allosterically by insulin-like growth factor 2. Sci Rep. 2024 Nov 6;14(1):26875. doi: 10.1038/s41598-024-75300-9. PMID: 39505925; PMCID: PMC11541866.

52. Kornfeld, S. Trafficking of lysosomal enzymes in normal and disease states. Journal of Clinical Investigation 77, 1–6 (1986).

53. Tong, P. Y., Tollefsen, S. E. & Kornfeld, S. The cation-independent mannose 6-phosphate receptor binds insulin-like growth factor II. J Biol Chem 263, 2585–8 (1988).

54. Alpern D, Gardeux V, Russeil J, Mangeat B, Meireles-Filho ACA, Breysse R, Hacker D, Deplancke B. BRB-seq: ultra-affordable high-throughput transcriptomics enabled by bulk RNA barcoding and sequencing. Genome Biol. 2019 Apr 19;20(1):71. doi: 10.1186/s13059-019-1671-x. PMID: 30999927; PMCID: PMC6474054.

