## Supplemental Figures/Tables for "Subtle Cellular Phenotypes Inform Pathological and Benign Genetic Mutants in the Iduronidase-2 Sulfatase Gene"

### Tables and Supplements

#### Supplemental Table 1

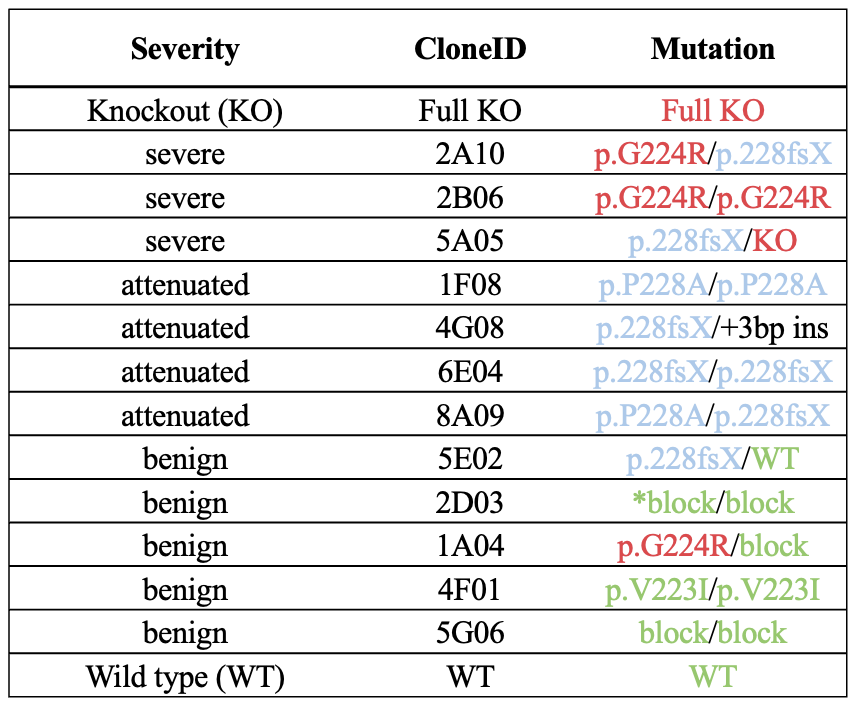

**Supplemental Table** 1. Details of the genotype of each cloneID, listing the combined genotypes and the associated classifications of those 12 selected variants. Red indicates a severe pathogenic mutant allele, blue is attenuated pathogenic, and green is benign.

#### Supplemental Table 2

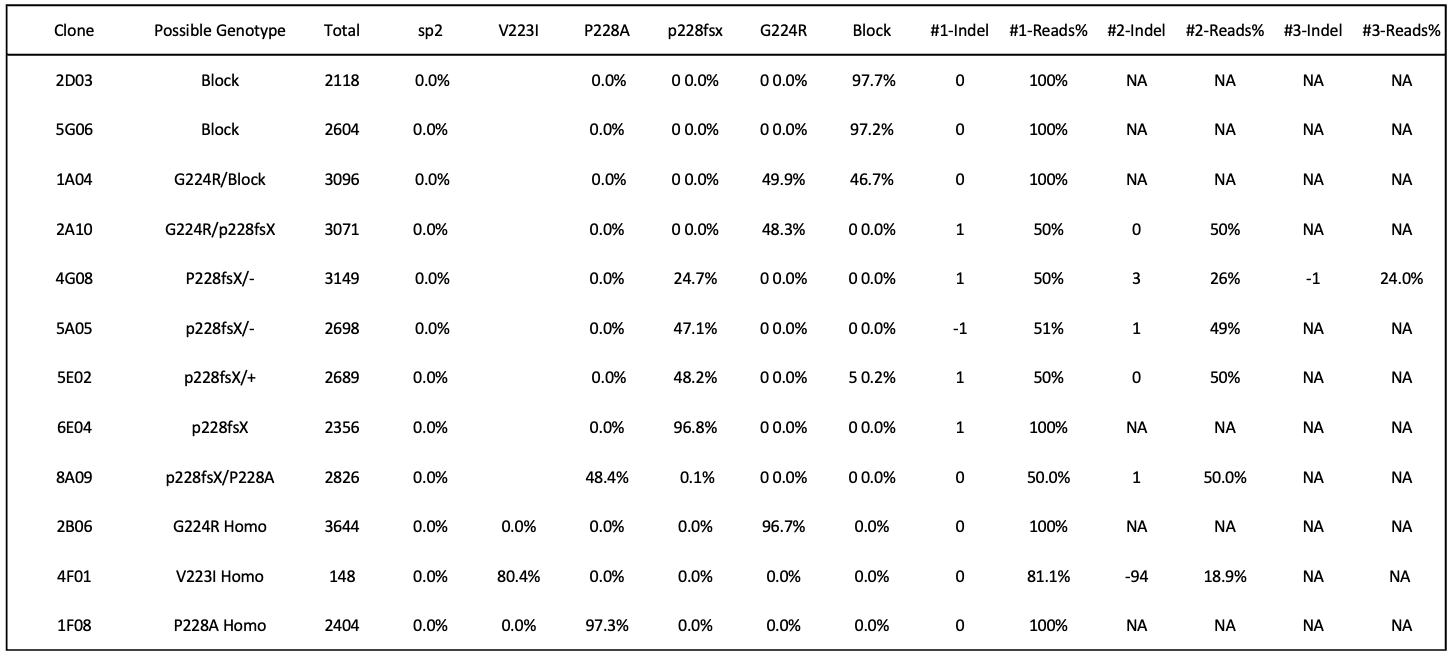

**Supplemental Table 2.** Clone information regarding the A549 IDS mutant lines. Each row shows a clone ID, and then the suspected genotype, and the Total number of reads. Each column with % in the values are indicating the percent of the reads that have the particular gRNA (sp2) or mutation V223I, P228A, p228fsx, G224R, Block present. Next is the most common (#1) indel, which is either a positive number (insertion), negative number (deletion), or 0 indicating no indel. Additional indels are listed but are usually not present.

#### Supplemental Table 3

| CELLS AREA WVC | CELLS KURTOSIS WVC | LYSOS TOTAL AREA WVL |
| --- | --- | --- |
| CELLS CHORD RATIO WVC | CELLS KURTOSIS WVL | LYSOS TOTAL INTENSITY WVL |
| CELLS COMPACTNESS WVC | CELLS LIGHT FLUX WVC | NUCLEI ENERGY WVC |
| CELLS DIAMETER WVC | CELLS LIGHT FLUX WVL | NUCLEI ENERGY WVL |
| CELLS ELONGATION WVC | CELLS LOW GREY LEVEL RE WVC | NUCLEI ENTROPY WVC |
| CELLS ENERGY WVC | CELLS LOW GREY LEVEL RE WVL | NUCLEI ENTROPY WVL |
| CELLS ENERGY WVL | CELLS MAX HEIGHT WVC | NUCLEI GREY LEVEL NU WVC |
| CELLS ENTROPY WVC | CELLS MAX INTENSITY WVC | NUCLEI GREY LEVEL NU WVL |
| CELLS ENTROPY WVL | CELLS MAX INTENSITY WVL | NUCLEI HIGH GREY LEVEL RE WVC |
| CELLS FORM FACTOR WVC | CELLS MAX WIDTH WVC | NUCLEI HIGH GREY LEVEL RE WVL |
| CELLS GREY LEVEL NU WVC | CELLS NUC/CELL AREA WVC | NUCLEI INTENSITY CV WVC |
| CELLS GREY LEVEL NU WVL | CELLS PERIMETER WVC | NUCLEI INTENSITY SD WVC |
| CELLS GYRATION RADIUS WVC | CELLS RUN LENGTH NU WVC | NUCLEI INTENSITY SD WVL |
| CELLS HIGH GREY LEVEL RE WVC | CELLS RUN LENGTH NU WVL | NUCLEI INTENSITY WVC |
| CELLS HIGH GREY LEVEL RE WVL | CELLS SKEWNESS WVC | NUCLEI INTENSITY WVL |
| CELLS INTENSITY (CELL) WVC | CELLS SKEWNESS WVL | NUCLEI KURTOSIS WVC |
| CELLS INTENSITY (CELL) WVL | CELLS TOTAL INTENSITY (CELL) WVC | NUCLEI KURTOSIS WVL |
| CELLS INTENSITY (CYTO) WVC | CELLS TOTAL INTENSITY (CELL) WVL | NUCLEI LIGHT FLUX WVC |
| CELLS INTENSITY (CYTO) WVL | CELLS TOTAL INTENSITY (CYTO) WVC | NUCLEI LIGHT FLUX WVL |
| CELLS INTENSITY CV (CELL) WVC | CELLS TOTAL INTENSITY (CYTO) WVL | NUCLEI LOW GREY LEVEL RE WVC |
| CELLS INTENSITY CV (CELL) WVL | LYSOS AREA WVL | NUCLEI LOW GREY LEVEL RE WVL |
| CELLS INTENSITY CV (CYTO) WVC | LYSOS DISTANCE TO NUC WVL | NUCLEI MAX INTENSITY WVC |
| CELLS INTENSITY CV (CYTO) WVL | LYSOS FORM FACTOR WVL | NUCLEI MAX INTENSITY WVL |
| CELLS INTENSITY SD (CELL) WVC | LYSOS INTENSITY SPREADING WVL | NUCLEI RUN LENGTH NU WVC |
| CELLS INTENSITY SD (CELL) WVL | LYSOS INTENSITY WVL | NUCLEI RUN LENGTH NU WVL |
| CELLS INTENSITY SD (CYTO) WVC | LYSOS NEIGHBOR COUNT WVL | NUCLEI SKEWNESS WVC |
| CELLS INTENSITY SD (CYTO) WVL | LYSOS ORG PER CELL WVL | NUCLEI SKEWNESS WVL |
| CELLS INTENSITY SPREADING WVC | LYSOS ORGANELLE/CYTO INTENSITY WVL | NUCLEI TOTAL INTENSITY WVC |
| CELLS INTENSITY SPREADING WVL | LYSOS SPACING WVL | NUCLEI TOTAL INTENSITY WVL |

**Supplemental Table 3.** A listing of all the imaging features that are included in the PathScore_LC_. The first word indicates the mask, so Nuclei means the pixels that are over the nuclei. The final “WV” phrase indicates the fluorescent channel that is measured, where C is membrane, L is lysosome, and H is nucleus. The middle words indicate the type of measurement that was made.

#### Supplemental Figure 1

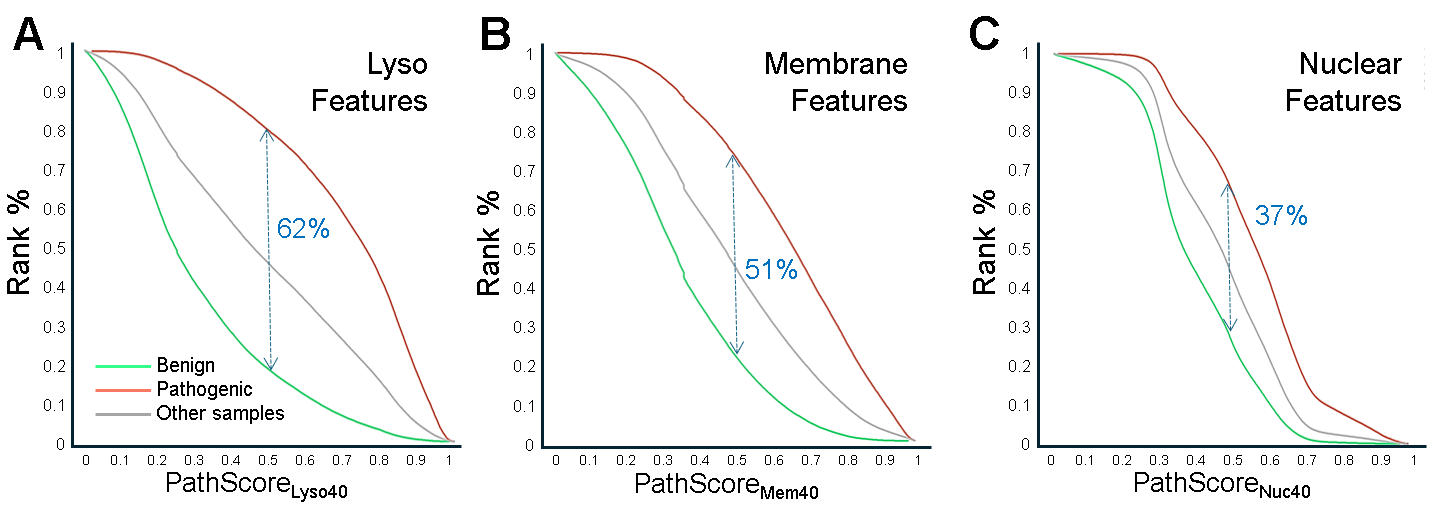

**Supplemental Figure 1.** Comparison of ability to separate Pathogenic and Benign Phenotypes with only **A**) Lysosomal features, **B**) Membrane features, and **C**) Nuclear features. Each figure represents the best model from a 22-set run, where there are 40 features sampled from the organelle specified. Each run had 2,500 Epochs, and Dense layers of size 54, 59, 80.

#### Supplemental Figure 2

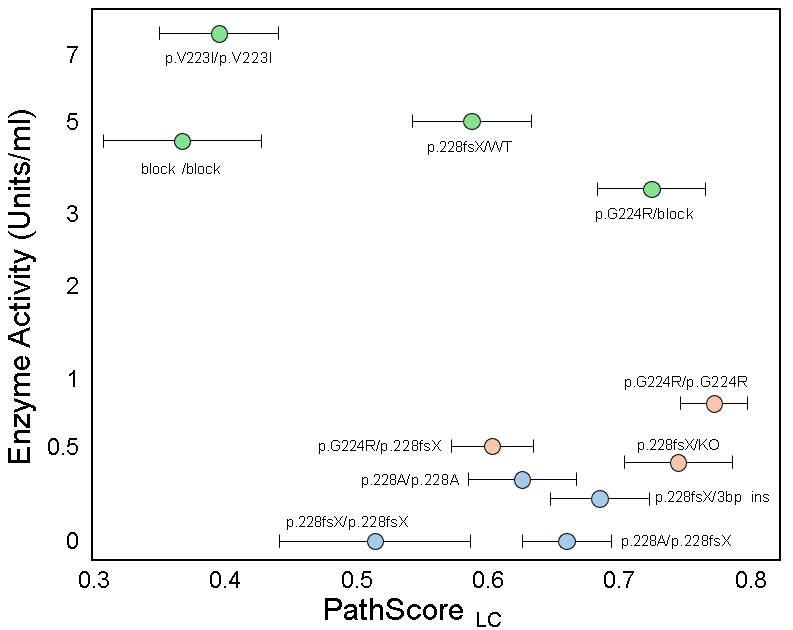

**Supplemental Figure S2.** Scatter plot showing the mean of all wells for each mutation-containing A549 cell line. Each point consists of at least 3 separate plates with 8 wells each and thousands of cells. Error bars are standard deviation. The mutation is indicated and plotted alongside the enzyme units.
